# Stoichiometry and Mobility Switching of a Morphogenetic Protein in Live Differentiating Cells

**DOI:** 10.1101/356733

**Authors:** Adam J. M. Wollman, Katarína Muchová, Zuzana Chromiková, Anthony J. Wilkinson, Imrich Barák, Mark C. Leake

## Abstract

Spore formation following asymmetric cell division in *Bacillus subtilis* offers a model system to study development, morphogenesis and signal transduction in more complex organisms. Extensive biochemical and genetic details of its sporulation factors are known, however, the molecular mechanisms by which asymmetry is generated remain unclear. A crucial membrane phosphatase, SpoIIE, couples gene regulation to morphology changes, but how it performs different functions dependent on cell stage is unknown. We addressed this puzzle using high-speed single-molecule fluorescence microscopy on live *B. subtilis* expressing genomically encoded SpoIIE fluorescent protein fusions during sporulation. Copy number analysis indicated a few tens of SpoIIE at sporulation onset increasing to 400-600 molecules per cell following asymmetric cell division with up to 30% greater proportion in the forespore, corresponding to a concentration enhancement in the smaller forespore sufficient for differential dephosphorylation of an anti-sigma factor antagonist and activation of the forespore specific transcription factor, σ^F^. Step-wise photobleach analysis indicates that SpoIIE forms tetramers capable of reversible oligomerisation to form clusters correlated with stage-specific functions. Specifically, low mobility SpoIIE clusters which initially localize to the asymmetric septum are released as mobile SpoIIE clusters around the forespore when phosphatase activity is manifested. SpoIIE is subsequently recaptured at the septum in a SpoIIQ-dependent manner. After mother cell engulfment of the forespore, SpoIIE is released as a mix of higher mobility clusters and tetramers. Our findings suggest that additional information captured in the changing state of multimerization and mobility enable one protein to perform different roles at different cell stages.

**Significance/impact:** Certain bacteria undergo sporulation involving cells dividing asymmetrically. A crucial protein SpoIIE facilitates this morphological asymmetry and directly links it to asymmetry in gene expression. Here, we used specialized light microscopy, capable of observing single molecules, plus biophysics, genetics and biochemical tools, to monitor SpoIIE in single living bacteria in real time, allowing us to count how many molecules are present in different cell regions, and how mobile they are. We find that SpoIIE clusters and moves depending on development stages, indicating that it has different roles depending on other binding proteins and their cellular locations. Our results suggest that changes in molecular stoichiometry and mobility may be used as switches in more complex cell processes.

---

Spore formation in *B. subtilis* offers a model system for studying development, differentiation, morphogenesis, gene expression and intercellular signalling in complex organisms(1, 2). In nutrient rich conditions, rod-shaped cells grow and multiply by symmetric mid-cell division to generate identical daughters (Fig. S1A). However, under starvation, asymmetric division produces a smaller forespore cell next to a larger mother cell. Each compartment inherits an identical chromosome, but patterns of gene expression, orchestrated by compartment-specific RNA polymerase sigma factors, differ resulting in alternative cell fates. The mother cell engulfs the forespore in a phagocytosis-like process creating a cell-within-a-cell (Fig. S1A), and a nurturing environment in which a robust multi-layered coat is assembled around the maturing spore(3). In the final stages, the mother cell undergoes programmed cell death releasing the spore, which is resistant to multiple environmental stresses and can lie dormant until favorable growth conditions are restored.

At sporulation onset, ring-like structures of the tubulin homologue FtsZ form at mid-cell and migrate on diverging spiral trajectories towards the cell poles(4), colocalizing with the membrane integrated phosphatase SpoIIE(5). One polar ring matures into the sporulation septum while the other disassembles(6). Asymmetric division otherwise involves the same proteins as vegetative cell division, though the resulting sporulation septum is thinner(7, 8). SpoIIE is the only sporulation-specific protein whose mutation causes ultrastructural changes in the asymmetric septum; null mutants of *spoIIE* are defective in sporulation and at low frequency give rise to thicker asymmetric septa resembling the vegetative septum(7).

Changes in cell morphology during sporulation are coupled to a programme of gene expression, involving intercellular signalling, and the sequential activation of RNA polymerase sigma factors, σ^F^ and σ^G^ in the forespore and σ^E^ and σ^K^ in the mother cell(9). Forespore-specific activation of σ^F^ on completion of the asymmetric septum is the defining step in differentiation. In pre-divisional and mother cells, σ^F^ resides in complex with the antisigma factor SpoIIAB while a third protein SpoIIAA is phosphorylated. Following SpoIIE activity on its substrate SpoIIAA~P, SpoIIAA displaces σ^F^ from the σ^F^:SpoIIAB complex allowing RNA polymerase binding and transcription of forespore-specific genes (Fig. S1B)(10, 11), triggering activation of σ^E^ in the mother cell and alternate programmes of gene expression which determine different cell fates (Fig. S1A).

SpoIIE has multiple roles at different sporulation stages (Fig. S1). Assembly of SpoIIE to form polar E-rings, dependent on FtsZ interactions (12), occurs during stage I, defined by formation of an axial filament spanning the cell length and comprising two copies of the chromosome each tethered through its origin region to a pole. Formation of the asymmetric septum is defined as stage II_i_, during which SpoIIE interacts with the divisome components RodZ(13) and DivIVA(14). After closure of the sporulation septum, FtsZ disassembles. SpoIIE-mediated activation of σ^F^ correlates with release of SpoIIE from the sporulation septum, marking stage II_ii_(13). During stage II_iii_, SpoIIE interacts with SpoIIQ(15) the forespore component of an intercellular channel(16–18), crucial for later activation of σ^G^. Stage III is characterized by mother cell engulfment of the forespore; SpoIIE localizes around the forespore, but there are no data to suggest a specific role of SpoIIE in this or later stages(15).

To clarify SpoIIE’s physiological roles, we sought to determine its dynamic molecular architecture and protein interactions. We employ high-speed Slimfield imaging(19–21) capable of tracking single fluorescently-labeled SpoIIE molecules with rapid millisecond sampling in live *B. subtilis* cells. By using step-wise photobleaching of the fluorescent protein tag(22) we determine the stoichiometry of each tracked SpoIIE complex. Also, by analysing their mean square displacement with respect to time we calculate the microscopic diffusion coefficient *D*, model this to determine the effective diameter of SpoIIE complexes, and correlate these data with measured SpoIIE content. We find that the stoichiometry and diffusion of tracked SpoIIE is dependent on its interaction partners and morphological changes suggesting its roles in sporulation are influenced by oligomeric composition and mobility. Interestingly, we distinguished higher order mobile, oligomeric SpoIIE at the stage of sporulation when σ^F^ becomes selectively activated in the forespore.

## Results

### SpoIIE copy number is 10-30% higher in early forespores

We generated a chromosomally encoded fusion of SpoIIE to monomeric yellow fluorescent protein mYPet (a bright fluorescent protein with short maturation time whose long excitation wavelength results in minimal contamination of cellular autofluorescence(23)) to report on SpoIIE localization (Table S1). We prepared cells for sporulation using nutrient-depleted media, incubating with red lipophilic dye FM4-64 for visualising *B. subtilis* membrane structures(24). This allowed us to observe steady-state patterns of SpoIIE-mYPet and FM4-64 localization for sporulation stages I, II (with associated sub-stages) and III with single-molecule precise Slimfield (Fig. 1A) and standard epifluorescence microscopies (Fig. 1B). We used bespoke single particle localization(25) on Slimfield data to track distinct SpoIIE foci whose width was consistent with the measured point spread function (PSF) of our microscope, ~250nm (Movies S1-3). Foci could be tracked over consecutive images up to ~0.3s using rapid 5ms per frame sampling to a spatial precision of 40nm(26). Generating frame averages over the first five Slimfield frames resulted in images qualitatively similar to those from epifluorescence (Fig. S2). We developed an automated high-throughput analysis framework using morphological transformations(27, 28) on SpoIIE-mYPet and FM4-64 data, enabling us to categorise each cell into one of five different sporulation stages (I, forespore formation stages II_i_, II_ii_ and II_iii_, and III after engulfment), validated by simulation and principal component analysis (Fig. 1C, Fig. S2 and SI Appendix). The measured proportions of cells in each stage (Fig. 1D) were qualitatively similar to those reported using manual, low-throughput methods(29). Imaging a SpoIIE-mYPet strain including a *ΔspoIIQ* deletion (Table S1), defective in spore formation and unable to progress beyond stage II_ii_, yielded similar relative proportions of cells in stages I, II_i_ and II_ii_ with some falsely categorised as stage III in low proportion compared to wild type (Fig. S2).

**Figure 1:**
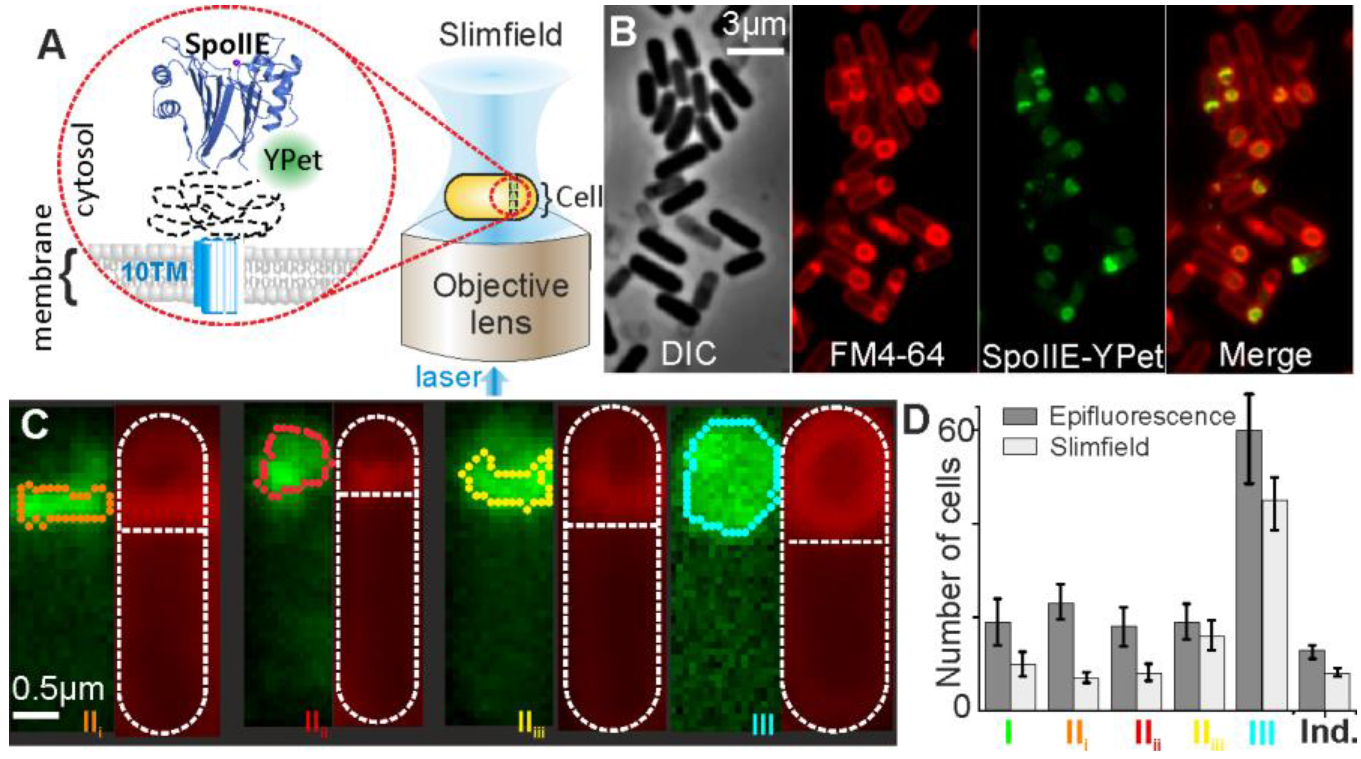
(A) Slimfield schematic, inset showing septum-localised SpoIIE-mYPet. (B) DIC/Epifluorescence of SpoIIE at different stages. Membrane labelling, FM4-64 (red), SpoIIE-YPet (green). (C) Categorization of stage from Slimfield, detected forespore/septa features (colored lines), cell boundary segmentation based on fitting a sausage shape to fluorescence image indicated (outer white dash) and interface between forespore and mother cell (horizontal white dash). (D) Detected mean frequency of each stage (SEM error) from N=4 fields of view each containing approximately N=100 cells.

Tracking was coupled to stoichiometry analysis by estimating the initial foci brightness and dividing this by the brightness of a single mYPet (Fig. S3). We also observed a diffuse pool of mYPet fluorescence (with negligible cellular autofluorescence) consistent with contributions from foci beyond our microscope’s depth of field (SI Appendix). By using integrated pixel intensities(26), we determined the total SpoIIE copy number for each cell. Utilizing our stage categorization algorithm we then quantified the number of SpoIIE specifically associated with the mother cell and the forespore (Fig. 2A, S4 and Table S2).

**Figure 2:**
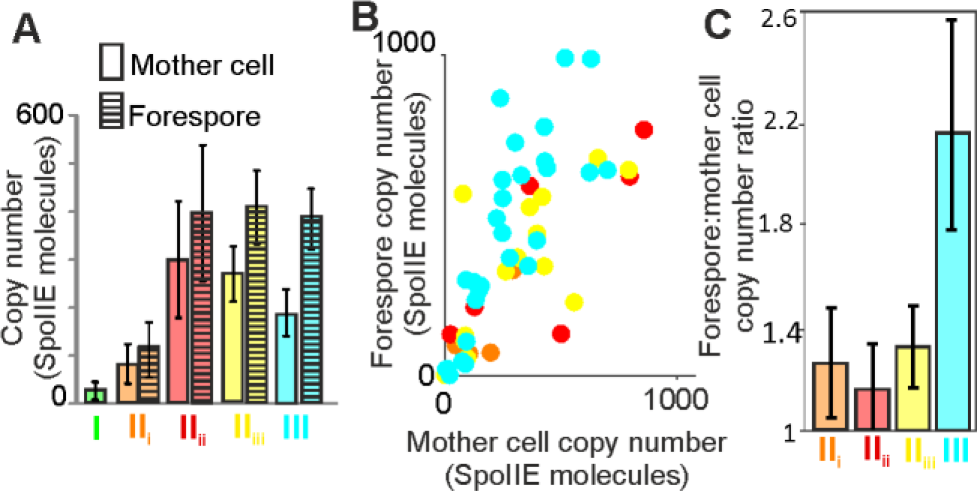
(A) Mean and SEM SpoIIE copy number in mother cell and forespore at each stage. (B) Scatter plot of forespore *vs.* mother cell copy number. (C) Mean and SEM ratio of forespore to mother cell copy number from linear fits of (B).

These analyses show the total SpoIIE copy number starts at a few tens of molecules per cell in stage I, increasing as sporulation progresses to ~200 SpoIIE in stage II_i_. SpoIIE copy number in the forespore plateaus at ~400 molecules at stages II_ii_-III but peaks at stage II_ii_ in the mother cell at ~300 copies before decreasing. Although *spoIIE* mRNA levels reach similar peak levels to those of *ftsZ*(30) the FtsZ copy number is estimated to be 2000-4000 molecules per cell(31), ~5-fold higher than we find for SpoIIE. This 5:1 ratio is very similar to FtsZ:FtsA in *B. subtilis(32)*, where it was suggested that FtsA forms a ring at the division site tethering FtsZ filaments to the membrane. We did not observe obvious patterns of spiral SpoIIE trajectories as had been reported in a previous study for FtsZ-GFP(4). We find that SpoIIE copy number in the forespore is directly proportional to, and 10-30% higher (Fig. 2B,C) than, the mother cell copy number in all stages preceding stage III when SpoIIE copy number decreases in the mother cell.

### SpoIIE is a tetramer in the mother cell independent of stage

In the mother cell, the apparent stoichiometry of tracked foci ranged from as few as 2 up to several tens of molecules, but with a clear peak at 4±2 SpoIIE molecules, conserved throughout stages I-III (Fig. 3). Using a randomized Poisson model for nearest-neighbor foci distances, comprising SpoIIE copy number and foci density, we calculated the probability of foci being separated by less than the optical resolution limit (thus detected as single foci of higher apparent stoichiometry) to be 20-40% in the mother cell. Overlap models which used SpoIIE monomers, dimers, hexamers or octamers do not account for the apparent stoichiometry data (Fig. S5 and SI Appendix). By contrast, we find that a tetramer overlap model generates reasonable agreement within experimental error for stages I-III in the mother cell (Fig. 3, dashed lines). Thus, SpoIIE in the mother cell comprises predominantly tetramers.

**Figure 3:**
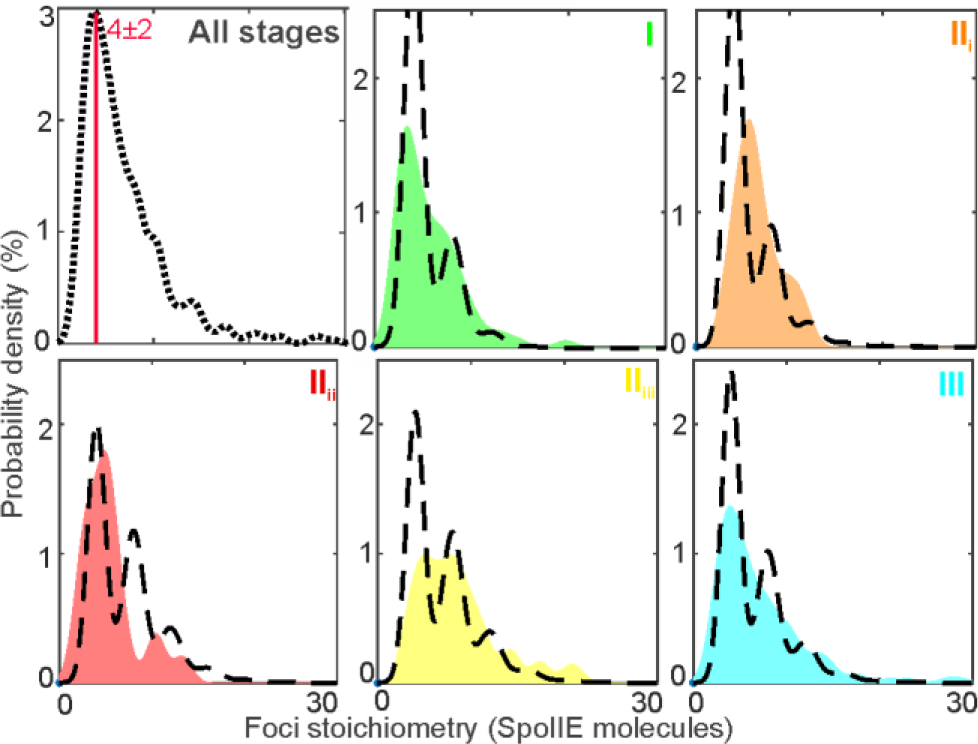
Kernel density estimation (KDE) of stoichiometry in mother cell with predicted overlap tetramer model (black dotted line). Goodness-of-fit *R*^*2*^ in range 0.3-0.7.

### Forespore SpoIIE is a tetramer which assembles into higher stoichiometry clusters according to stage

For forespore SpoIIE foci, we find the same tetramer peak in the apparent stoichiometry distribution but with a longer tail of higher stoichiometry clusters extending up to hundreds of SpoIIE molecules per foci (Fig. 4A, B). We adapted the overlap model to account for different sizes and shapes of sporulation features at each stage resulting in differences in the density of SpoIIE foci (including a septum at stages II_i_ and II_iii_ and a complete forespore in II_ii_ and III) (Fig. 4B). The overlapping tetramer model accounts only for low apparent stoichiometries near the tetramer peak and only in stages II_iii_ and III. More generally, to account for the apparent stoichiometry in the forespore requires populations of higher order oligomeric SpoIIE clusters in addition to tetramers. Excluding free tetrameric foci, we observe 1-3 clusters per cell (Fig. 4D) with the mean cluster stoichiometry peaking in stage II_ii_ at >100 (Fig 4C) before decreasing as the proportion of free tetramers increases (Fig, S7A, B). We find that forespore foci apparent stoichiometry in all stages is periodic around 4 molecules (Fig. S6), suggesting that higher order clusters are composed of associating SpoIIE tetramers.

**Figure 4:**
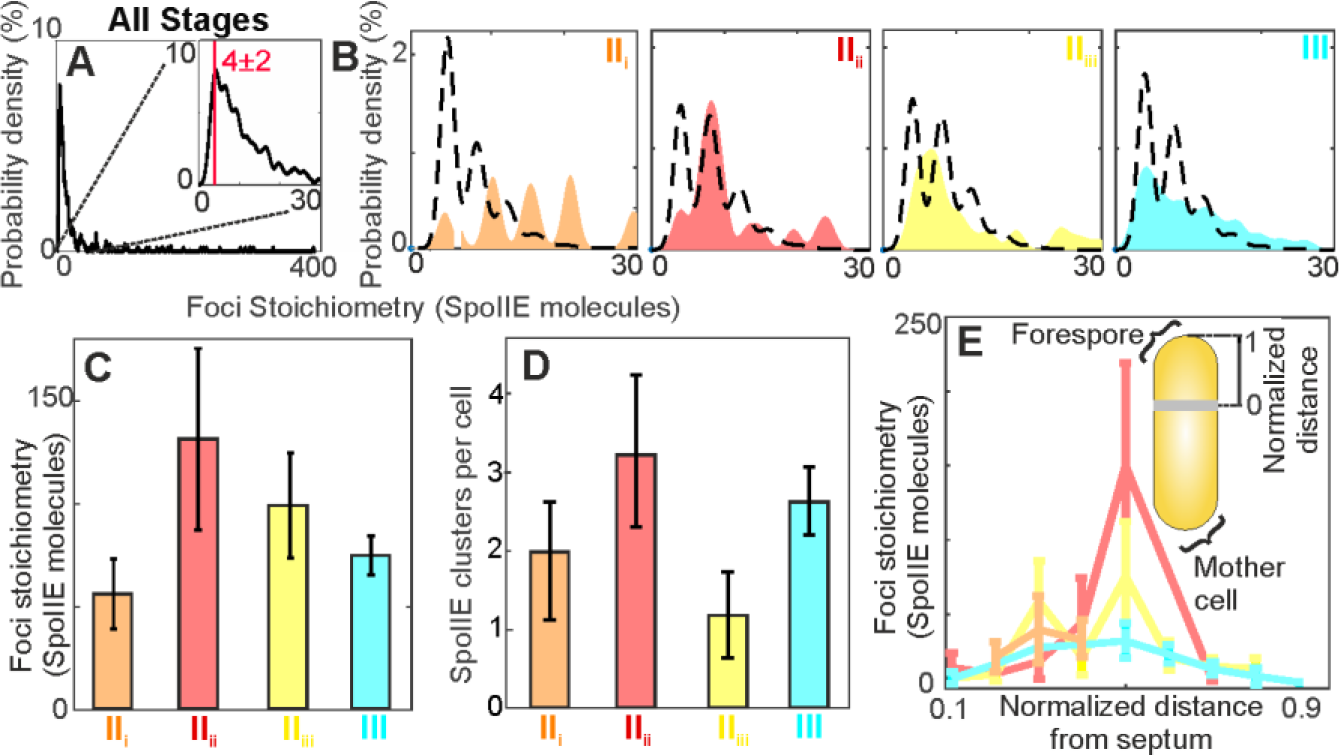
(A) KDE of forespore stoichiometry pooled for all data, and zoom-in (inset), (B) in separate stages with overlap tetramer model (black dotted line). (C) Bar chart of mean foci stoichiometry for foci which have >20 molecules. (D) Mean number of foci detected per forespore. (E) Mean foci stoichiometry *vs.* normalised distance from septa for each stage, (inset) schematic of forespore distance normalisation.

We also find spatial dependence of foci stoichiometry in the forespore. We normalised the distance parallel to the long axis of each cell from the mother cell side of the asymmetric septum through to the outer edge of the cell for all tracked foci (Fig. 4E inset) and plotted this against stoichiometry (Fig. S7C and SI Appendix). For stage II_i_, foci are localized to a region ~400nm from the septum itself, however, other stages contain foci which are delocalized over the full extent of the emerging forespore (Fig. 4E); we find an increase in mean SpoIIE stoichiometry for these by a factor of ~12 for stage II_ii_ with a smaller ~1.4-fold increase in stage II_iii_.

In stage II_i_, when SpoIIE is likely to be associated with the divisome, we observe ~2 clusters per cell, with a mean stoichiometry of ~50. When SpoIIE is released from the divisome complex in stage II_ii_, we find ~3 clusters per cell with a much higher stoichiometry of ~130 molecules per foci, relatively far from the septum in the direction of the pole (Fig. 4E). Clustering of SpoIIE in the direction of the pole was recently proposed as a mechanism for σ^F^ activation regulation(33). As SpoIIE is recaptured at the septum by interaction with SpoIIQ(15), we find that the high polar stoichiometry decreases, as does the mean cluster stoichiometry, and only one cluster is observed together with a population of free tetramers. Interestingly, in stage III, when SpoIIE has no attributed function in sporulation, free tetramers are observed together with 2-3 clusters per cell.

### SpoIIE mobility depends on stage

We find that foci mobility in general is consistent with Brownian (i.e. normal) diffusion irrespective of cell compartment or stage (Fig. S8). In the mother cell, the mean *D* is 0.9-1.2μm^2^/s while that in the forespore is lower by a factor of ~2 (Fig. 5A, Fig. S8 and Table S2). At the onset of sporulation in stage II_i_ foci mobility in the forespore is at its lowest with a mean *D* of 0.43±0.08μm^2^/s, which increases during stage II_ii_ to 0.67±0.19μm^2^/s, then decreases in stage II_iii_ to 0.50±0.09μm^2^/s before increasing again in stage III to 0. 76±0.05μm^2^/s. For stages I-III, *D* shows a dependence on stoichiometry *S*, indicating a trend for decreasing *D* with increasing SpoIIE content (Fig. 5B). Modeling this dependence as *D* ~ *S^α^* indicates a power-law exponent α of 0.48 ± 0.18, with no measurable difference within error for each stage (Fig. S9 and S10).

**Figure 5:**
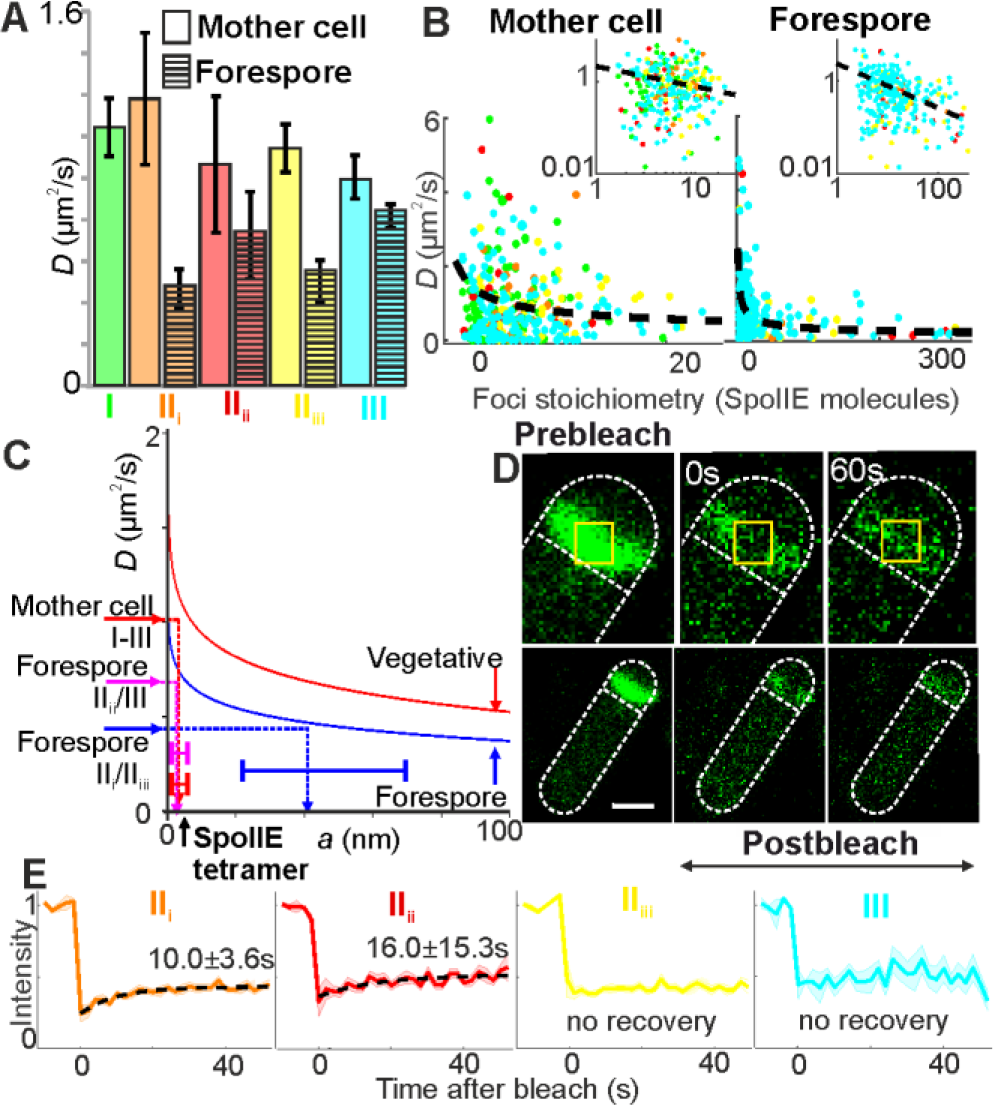
(A) Bar chart of mean *D* for each compartment and stage, SEM errors. (B) Scatter plots of stoichiometry *vs. D* for all stages in mother cell and forespore, log-log axes inset (power law model black line). (C) Variation of *D* with effective cluster cylinder radius *a* from frictional drag model, vegetative (red) and forespore (blue) indicated with interpolations from *D* to *a* made from mother cell (all stages), and forespore stages (II_i_/II_iii_) and (II_ii_/III), SEM errors. (D) FRAP from representative cell in stage II_i_, immediately pre/post-bleach, and ~60s post-bleach, bleached region indicated (yellow squares). (E) Mean normalised fluorescence recovery for each stage, SEM bounds shown (shading). Dotted lines show best exponential fit where recovery was detected, *t*_*r*_ and 95% confidence intervals indicated.

We also identify *D* as equal to *k*_*B*_*T*/γ using the Stokes-Einstein relation, where *k_B_* is the Boltzmann constant and *T* the absolute temperature, modeling lateral viscous drag *γ* on SpoIIE foci as predominantly due to friction between the phospholipid bilayer and SpoIIE transmembrane regions since the viscosity in the plasma membrane is higher by 2-3 orders of magnitude than in the cytoplasm(34). We use a generalized model, characterizing transmembrane protein regions as cylinders to account for rotation in the plane normal to the membrane (35, 36), applying previous estimates for the relatively low and high viscosities of vegetative and forespore membranes respectively(37) (SI Appendix). Calculations using a consensus value for *D* from stages I-III for the mother cell indicate an average Stokes radius (the radius of equivalent cylinder in the membrane) in the range 3-8nm (Fig. 5C, red dashed line). Structural data predict 10 transmembrane alpha helices per SpoIIE subunit (38) each of diameter 1.2nm, thus a SpoIIE tetramer comprising 40 transmembrane helices has a ~4nm radius assuming close packing of helices, consistent with our experimentally-derived estimate. By contrast, a ‘mean’ ~50-mer SpoIIE cluster has a Stokes radius of ~13nm.

For the forespore, the average *D* for higher SpoIIE mobility stages II_ii_ and III indicates a similar range for Stokes radius to that calculated for foci in the mother cell (Fig. 5C, magenta dashed line). However, the low SpoIIE mobility stages II_i_ and II_iii_ indicate a Stokes radius approximately an order of magnitude higher at ~40nm (Fig. 5C, blue dashed line), far more than expected for a cluster of 100 SpoIIE molecules. This observation supports a model in which SpoIIE interacts with other proteins or complexes. In stage II_i_ interactions would be with components of the divisome(12–14) while in stage II_iii_ they would be with SpoIIQ, the forespore component of an intercellular channel formed with proteins encoded on the *spoIIIA* operon expressed in the mother cell(15).

Using confocal microscopy we performed fluorescence recovery after photobleaching (FRAP) experiments to photobleach the asymmetric septum at different stages and monitor any subsequent fluorescence recovery (Fig. 5D). During stages II_i_ and II_ii_ there is a relatively slow recovery with mean exponential recovery time *t*_*r*_ of 10±3.6s and 16.0± 15.3s respectively (Fig. 5E, S11). Our finding that *t*_*r*_ is not directly correlated to *D* in each stage suggests that turnover here is reaction- as opposed to diffusion-limited; it may be limited by an effective off-rate as observed in other complex bacterial structures such as components of the flagellar motor(22, 39). In subsequent stages II_iii_ and III, no recovery is detectable within error, though lower levels of fluorescence and numbers of cells in stage III result in higher measurement noise which limits the sensitivity for detecting low levels of putative recovery. Our findings suggest that interactions between SpoIIE and other structures (such as the divisome, or the SpoIIQ-SpoIIIAH channel present from stage II_iii_) may be stronger in stages II_iii_ and III compared to II_i_ and II_ii_, thus prohibiting measurable turnover during the time scale of our FRAP measurements.

## Discussion

SpoIIE performs multiple vital functions but how it switches roles at different stages has been unclear. It is not known how SpoIIE localizes to the polar septum, how it causes FtsZ to relocalize from mid-cell to a pole, what role it plays in septal thinning, or how its SpoIIAA-P phosphatase activity is controlled so that σ^F^ activation is delayed until the asymmetric septum is completed(7, 11). How SpoIIE brings about forespore-specific activation of σ^F^ is a subject of particular interest(41). Plausible suggested mechanisms include preferential SpoIIE localization on the forespore face of the septum(42), transient gene asymmetry leading to accumulation of a SpoIIE inhibitor in the mother cell(41), and the volume difference between compartments leading to higher specific activity of equipartitioned SpoIIE(44, 45). Most recently, it was shown that mother cell restricted intracellular proteolysis of SpoIIE by the membrane bound protease FtsH is important for compartment-specific activation of σ^F^(33). Our findings indicate that SpoIIE operates as an oligomer whose stoichiometry and mobility switch in the forespore according to specific sporulation stage. In particular, complexes comprising four SpoIIE molecules predominate in the mother cell and at multiple stages in the forespore. Crucially, we observe reversible assembly of these tetrameric SpoIIE entities into higher order multimers during stage II_ii_ when the protein localises towards the pole and its latent protein phosphatase activity is manifested(33).

Unlike previous microscopy of YFP-labelled SpoIIE which suggested a pattern of localization almost exclusively in the forespore following asymmetric septation(33), our improved sensitivity shows SpoIIE content is only 10-30% higher in the early forespore compared to the mother cell. However, the smaller forespore volume results in a higher SpoIIE concentration by a factor of ~10. It was shown previously that a 10-fold difference in SpoIIE phosphatase activity towards its substrate SpoIIAA~P could account for all-or-nothing compartmental regulation of σ^F^ activity(46). The bias towards higher copy number values in the forespore aligns with the recent suggestion that SpoIIE captured at the forespore pole is protected against proteolysis(33). In this model, SpoIIE sequestered in the polar divisome, is handed-off to the adjacent forespore pole following cytokinesis: SpoIIE is protected from FtsH-mediated proteolysis due to oligomerisation, which is clearly described by our observations. Compartment specificity results from the proximity of the forespore pole to the site of asymmetric division.

Crystallographic and biophysical studies reveal that SpoIIE(590-827), comprising the phosphatase domain, is a monomer while SpoIIE(457-827), comprising the phosphatase plus part of the upstream regulatory domain, is dimeric (Fig. 6A)(47, 48). Comparison of these structures and mapping of mutational data onto them led to the proposal that PP2C domains in SpoIIE(590-827) and SpoIIE(457-827) crystals represent inactive and active states respectively. Activation is accompanied by a 45° rigid-body rotation of two ‘switch’ helices(48). This switch is set by a long α-helix in the regulatory domain which mediates dimerization (Fig. 6A). Movement of the switch helices upon dimerization translates a conserved glycine (Gly629 in SpoIIE) into the active site where it can participate in cooperative binding to two catalytic manganese ions. These ions are conserved in PP2C phosphatases and here would be expected to activate a water molecule for nucleophilic attack at the phosphorus of the phoshorylated serine 57 residue in SpoIIAA-P. The increase in SpoIIE stoichiometry observed upon activation *in vivo* is consistent with these structural findings although clearly larger assemblies are implied. Our data suggest these arise from further homomeric quaternary interactions mediated by the membrane binding domain and/or the component of the regulatory domain which has yet to be fully characterized. The results are consistent with the hand-off model(33) in which release from the divisome allows SpoIIE tetramers to diffuse away from the septum and self-associate to form high stoichiometry clusters in a spontaneous process with similarities to that observed for the plasmalemmal protein syntaxin(49). We speculate that the free energy of reassembly is used to flip the helical switch, allowing manganese acquisition and activation of phosphatase activity.

**Figure 6:**
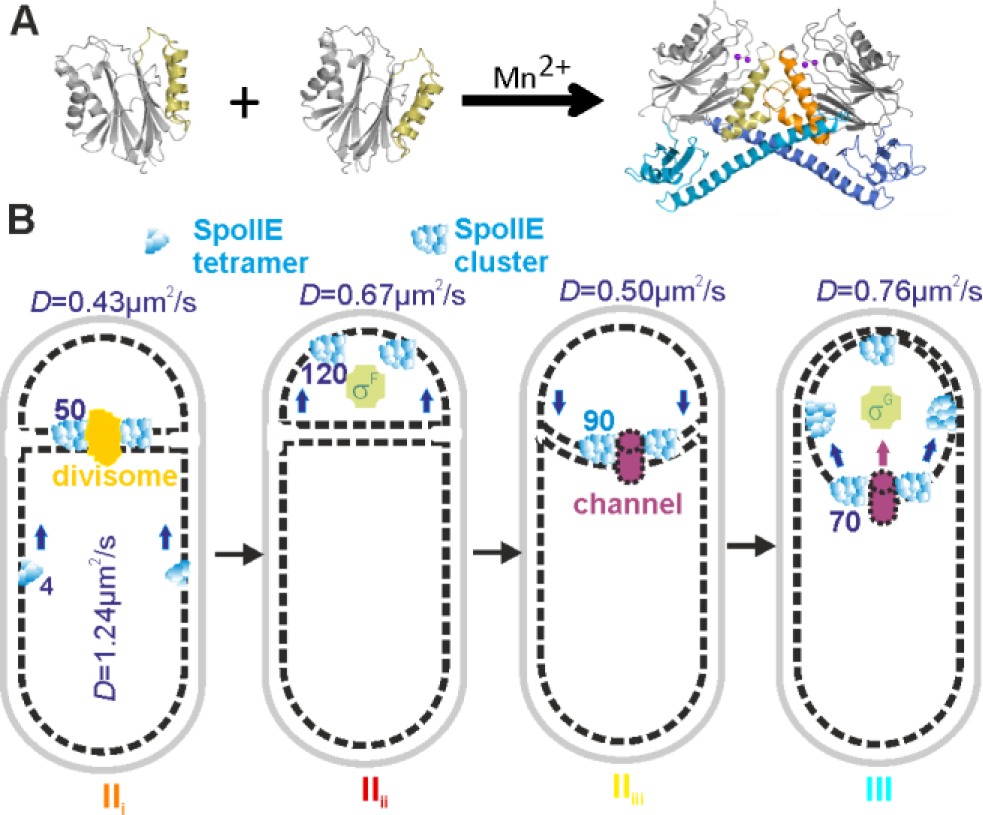
Model of SpoIIE dynamics during sporulation. (A) Activation of phosphatase involves dimerization, and recruitment of Mn^2+^ ions. Isolated phosphatase, SpoIIE(590-827) (PDB: 5MQH), is monomeric while a longer fragment, SpoIIE(457-827) (PDB: 5UCG), forms dimers about an interface dominated by a long helix spanning residues 473-518. Subunits in dimer distinguished by shading: PP2C domains (gray), switch helices (orange), regulatory domains (blue), Mn^2+^ ions (purple spheres, modeled onto structure following superposition of PstP structure from *M. tuberculosis* PDB: 1TXO). (B) Schematic of SpoIIE architecture and *D* at each stage: SpoIIE tetramers (blue), divisome (yellow), activated sigma factors (green), and SpoIIQ channel (purple) shown.

Changes in oligomeric state and quaternary organisation of proteins are widespread mechanisms for regulating biological activity. These can be induced variously by binding of allosteric ligands, covalent modification, proteolytic processing and reversible interactions with protein agonists or antagonists. SpoIIE, which transitions between complexes which are unusually large, is transiently active as a phosphatase after its release from an inhibitory complex with the divisome and prior to its recapture by SpoIIQ into an assembly that constitutes an intercellular channel. Regulation of phosphatase activity through sequestration is also seen in adaptation to drought in plants; the phosphatase HAB1 dephosphorylates the kinase SnRK2, inhibiting transcription of drought tolerance genes until the complex of the hormone, abscisic acid and its receptor, PYR, binds to and inhibits the phosphatase HAB1(50).

Our findings show more generally that we can combine robust cell categorization with single-molecule microscopy and quantitative copy number and stoichiometry analysis to follow complex morphologies during differentiation (Fig. 6B). Importantly, these tools provide new insight into the role of SpoIIE by monitoring its molecular composition and spatiotemporal dynamics, linking together different stages of cell development. Our findings show that the function of a key regulatory protein can be altered depending upon its state of multimerization and mobility, enabling different roles at different cell stages. Future applications of these methods may involve multicolor observations of SpoIIE with other interaction partners at different sporulation stages. Optimising these advanced imaging tools in the model Gram-positive *B. subtilis* may ultimately enable real time observations of more complex cellular development, paving the way for future studies of tissue morphogenesis in more challenging multicellular organisms.

## Materials and Methods

### Strains and plasmids

*B. subtilis* cloning followed standard protocols(53) (Table S1). Cells grown in DSM supplemented with antibiotics as required, were spotted on 1% agarose slides for microscopy 2h after sporulation onset. Full details SI Appendix.

### Microscopy

A Slimfield microscope was used(22, 27) for single-molecule imaging, an Olympus BX63 microscope measured epifluorescence. Tracking used MATLAB (Mathworks) software which determined *D* and stoichiometry using foci brightness(54) and Chung-Kennedy(55) filtered mYPet step-wise photobleaching, and nearest-neighbor modeling(56). FRAP was carried out on a Zeiss LSM 510 Meta confocal system. Full details, including cell categorization analysis, SI Appendix.

### Structural and bioinformatics analysis

CCP4mg was used to render images of structures with PDB IDs: 5MQH SpoIIE(590-827) and 5UCG SpoIIE(457-827) (57).

### Data availability

Data included in full in main text and supplementary files. Raw data available from authors.

### Software access

Code written in MATLAB available from Sporulationanalyser at https://sourceforge.net/projects/york-biophysics/.

## Acknowledgments

Authors thank Niels Bradshaw and Adam Hughes for helpful comments. Supported by Royal Society, MRC (grant MR/K01580X/1), BBSRC (grant BB/N006453/1) to M.L. and grants 2/0007/17 from Slovak Academy of Sciences and Slovak Research and Development Agency (APVV-14-0181) to I.B. Part-funded by the Wellcome Trust [ref: 204829] through Centre for Future Health at the University of York to A.J.M.W.

## Supplementary Appendix

### Supporting Tables

**Table S1.**
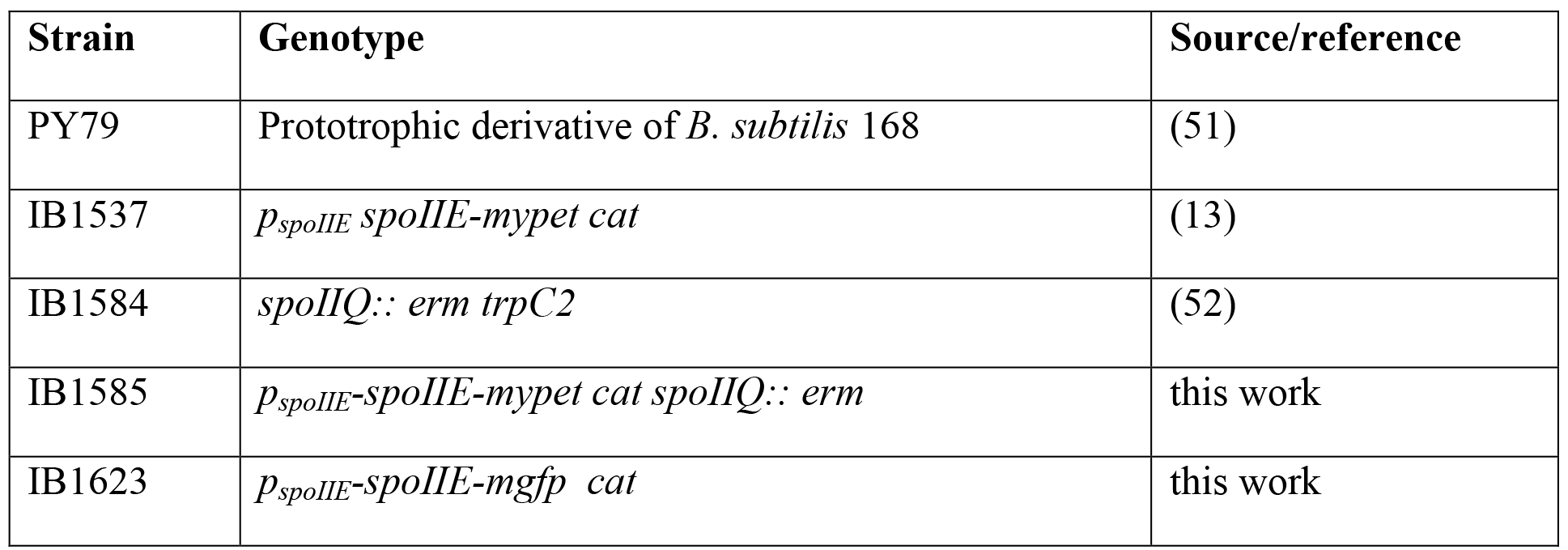
List of strains used in this study.

**Table S2.**
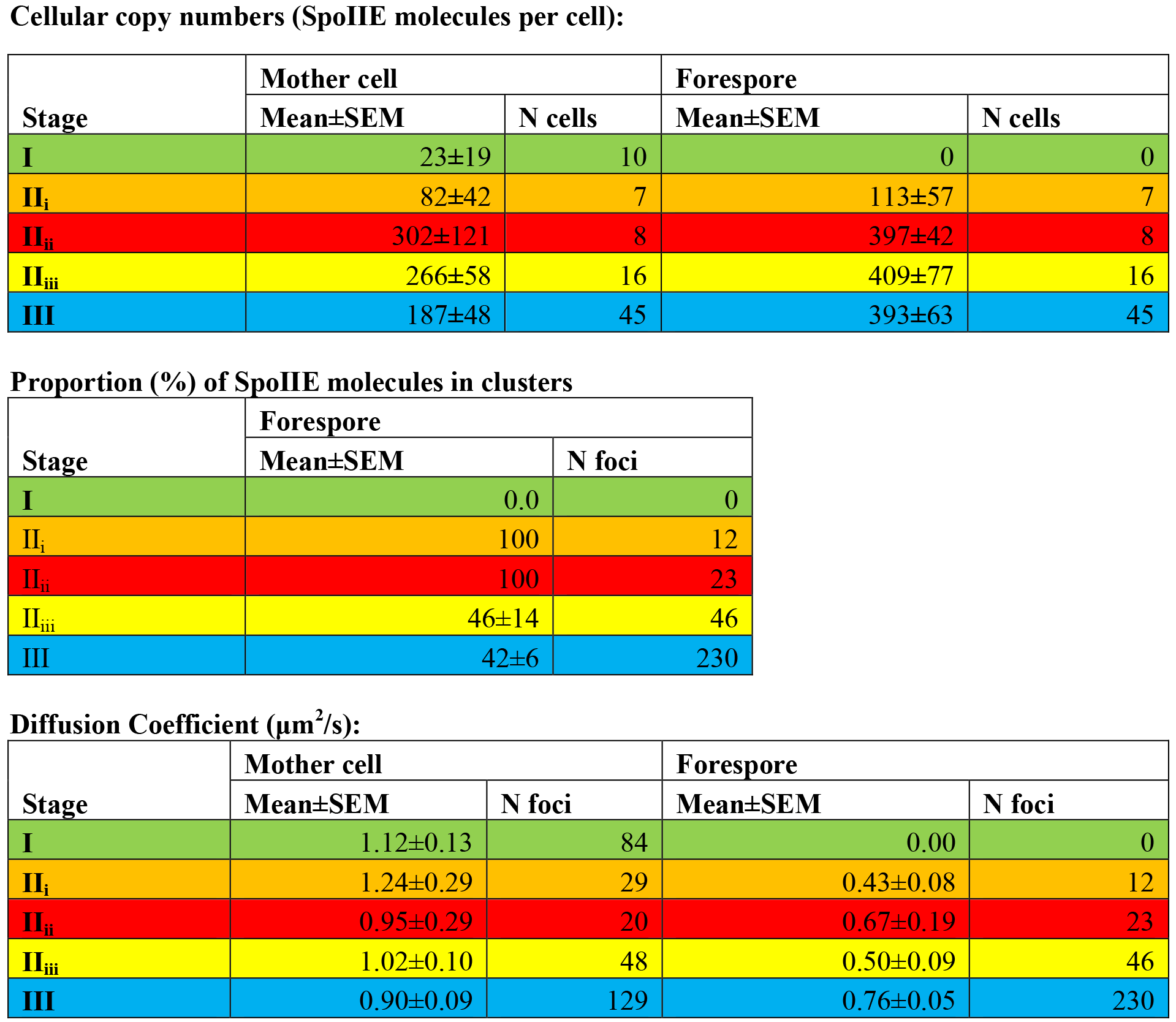
Summary of mean cellular copy numbers, foci stoichiometry and diffusion coefficient for WT SpoIIE-YPet. Number of cells (N cells) and foci (N foci) indicated, with SEM errors. Summary of mean cellular copy numbers, proportion of foci in clusters, and diffusion coefficients for WT SpoIIE-YPet foci. Number of cells (N cells) and foci (N foci) indicated, with SEM errors. In stages II_i_ and II_ii_ the stoichiometry distribution for foci cannot be account for by an overlapping tetramer and so by definition all foci are present in clusters.

**Cellular copy numbers (SpoIIE molecules per cell):**
| Stage | Mother cell |  | Forespore |  |
| --- | --- | --- | --- | --- |
|  | Mean±SEM | N cells | Mean±SEM | N cells |
| **I** | 23±19 | 10 | 0 | 0 |
| **II<sub>i</sub>** | 82±42 | 7 | 113±57 | 7 |
| **II<sub>ii</sub>** | 302±121 | 8 | 397±42 | 8 |
| **II<sub>iii</sub>** | 266±58 | 16 | 409±77 | 16 |
| **III** | 187±48 | 45 | 393±63 | 45 |

**Proportion (%) of SpoIIE molecules in clusters**
| Stage | Forespore |  |
| --- | --- | --- |
|  | Mean±SEM | N foci |
| **I** | 0.0 | 0 |
| **II<sub>i</sub>** | 100 | 12 |
| **II<sub>ii</sub>** | 100 | 23 |
| **II<sub>iii</sub>** | 46±14 | 46 |
| **III** | 42±6 | 230 |

**Table S2** Summary of mean cellular copy numbers, foci stoichiometry and diffusion coefficient for WT SpoIIE-YPet.
| Stage | Mother cell |  | Forespore |  |
| --- | --- | --- | --- | --- |
|  | Mean±SEM | N foci | Mean±SEM | N foci |
| **I** | 1.12±0.13 | 84 | 0.00 | 0 |
| **II<sub>i</sub>** | 1.24±0.29 | 29 | 0.43±0.08 | 12 |
| **II<sub>ii</sub>** | 0.95±0.29 | 20 | 0.67±0.19 | 23 |
| **II<sub>iii</sub>** | 1.02±0.10 | 48 | 0.50±0.09 | 46 |
| **III** | 0.90±0.09 | 129 | 0.76±0.05 | 230 |

## SI Appendix

### Strains and plasmids

Gene cloning in *B. subtilis,* unless specified, was performed using standard protocols (53) (Table S1). To construct pSGIIE-mGFP, we used previously prepared pSGIIE-YPet (13). A PCR fragment containing mGFP was prepared using mGFPKpnF: 5’ ATCATCATC<u>GGTACC</u>ATGAGTAAAGGAGAAGAAC TTTTCACTGGAGTTGTC 3’ and mGFPBamR2: 5’ atcatcatc<u>ggatcc</u>TTATTTGTATAGTTCATCCATGCCATGTG 3’a

primers and a plasmid derivative pSG1729 containing *mgfp* as template (58). To yield pSG-mGFP this fragment was KpnI/BamHI digested and cloned into a similarly cut pSGIIE-YPet. Subsequently a 360bp KpnI fragment containing *spoIIE* C-terminus (obtained from KpnI/BamHI cut pSGIIE-YPet) was cloned into pSG-mGFP digested with KpnI to yield pSGIIE-mGFP.

*B. subtilis* liquid cultures were grown in DSM (53) supplemented with chloramphenicol (5μg ml^−1^), erythromycin (1μg ml^−1^) and lincomycin (25μg ml^−1^) as required. Samples for microscopy on 1% agarose slides were taken 2h after sporulation onset. For membrane visualization, FM 4-64 (Molecular Probes) was used (0.2-1pg ml^−1^). When necessary, cells were concentrated by centrifugation (3min, 2,300x g) and resuspended in a small volume of supernatant. Images and analysis were obtained with an Olympus BX63 microscope (Hamamatsu Orca-R^2^ camera) and Olympus CellP or Olympus Image-Pro Plus 6.0 software.

### Single-molecule microscopy

A dual-color bespoke single-molecule microscope was used as described previously (21, 26) which utilized narrow epifluorescence excitation of 10μm full width at half maximum in the sample plane from a 514nm 20mW laser (Obis LS, Coherent). The laser was propagated through a ~3x Keplerian beam de-expander. Illumination was directed onto an *xyz* nanostage (Mad City Labs, the Dane County, Wisconsin, USA), and emissions directed through a color splitter utilizing a dichroic mirror centered on 560nm wavelength and emission 25nm bandwidth filters centered at 542/594nm (Chroma Technology Corp., Rockingham, Vermont, USA) onto an Andor iXon 128 emCCD camera, 80nm/pixel. Brightfield imaging was performed with no gain (100ms/frame), single-molecule imaging at maximum gain (5ms/frame).

Foci were automatically detected using MATLAB (Mathworks) software enabling a spatial localization precision of 40nm using iterative Gaussian masking, and automated *D* and stoichiometry calculation. The copy number in the mother cell or forespore was determined by summing pixel intensities within the compartment, correcting for low background autofluorescence measured from FM4-64 labeled wild type *B. subtilis*, then dividing by the characteristic SpoIIE-YPet intensity (26). The intensity of each foci was defined as the summed intensity inside a 5 pixel radius circle corrected for the local background, defined as the mean intensity in a 17 pixel square outside the circle (54). If the signal to noise ratio of the foci, defined as the mean intensity divided by the standard deviation of the local background, was greater than 0.4 it was linked into an existing track if within 5 pixels. The characteristic SpoIIE-YPet intensity was calculated from foci intensities found towards the end of the photobleach, confirmed to be single molecule from detection of single step-wise photobleach events in individual over-tracked, Chung-Kennedy(55) filtered SpoIIE-YPet tracks (Fig. S4). The stoichiometry of tracked foci was determined by fitting the first 4 intensity values of each track with exponential:

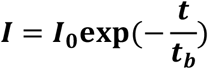

*I*=foci intensity, *I*_*0*_=initial intensity, *t*=time since laser illuminated cell, *t*_*b*_=bleach time (determined by an exponential fit to all population foci intensity to be ~100ms). *I*_*0*_ was divided by the YPet characteristic intensity to give the stoichiometry.

The 2D mean square displacement (MSD) was calculated from a fitted foci centroid (*x*(*t*),*y*(*t*)) assuming a track of *N* consecutive frames, and a time interval τ = *n*Δ*t*, where *n* is a positive integer and Δ*t* the frame integration time (59):

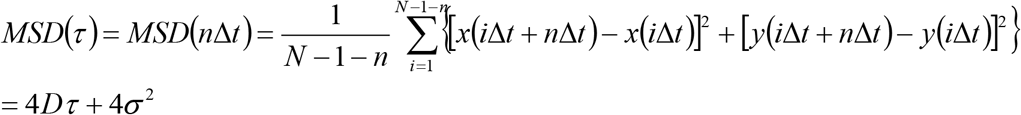

The localisation precision from tracking is given by *σ* which we measure as 40nm. *D* is estimated from a linear fit to the first three data points in the MSD *vs*. τ relation (i.e. 1 ≤ *n* ≤ 3) for each accepted track, with the fit constrained to pass through a point 4*σ*^2^ on the vertical axis corresponding to τ = 0, allowing *σ* to vary in the range 20 - 60nm in line with the experimental range.

### FRAP

FRAP was carried out on a Zeiss LSM 510 Meta confocal system with Axiovert inverted microscope, fitted with Plan Apochromat 100x /1.4 NA oil objective and temperature-controlled stage. A 488nm wavelength laser excited GFP, emissions collected via a 498-564nm bandpass filter. The strength of photobleaching in the region of interest was set to 10-20 iterations of 100ms each to ensure maximal photobleaching of GFP inside and minimum photobleaching beyond.

#### Categorization of cell cycle stage

To determine the cell cycle stage during the sporulation process, the following algorithm was used:
1. Cell images were initially coarsely over-segmented by thresholding the brightfield image and then using an initial ellipse shape approximation to define the cell length (26). We then manually optimised the cell width of a sausage function (rectangle capped with two hemicircles) that enclosed the YPet fluorescence intensity in each cell above the level of background noise.
2. Cells were then cropped out of the original image using a bounding rectangle around the segmentation and automatically rotated parallel to the horizontal axis.
3. A more precise segmentation stage then followed. This consisted of a double threshold Otsu’s method, applied to a 5 frame average of the YPet fluorescence image. Pixels whose intensity values were above the 2^nd^ threshold and multiplied by the segmentation contain the spore feature – either the whole forespore or septa.
4. These pixel areas were split into distinct connected components or candidate spore features and their centroids and areas calculated automatically using standard MATLAB functions.
5. A region was accepted as the YPet spore feature mask if:

1. Its centroid is within 40% of either end of the cell.
2. Its centroid is within ±40% of the middle of the cell width.
3. The area of its centroid was >10 pixels (there was no upper threshold).
4. It had the highest summed pixel intensity of all the regions.
6. If nothing was accepted, steps 5.1-5.4 were repeated once with the previously found regions excluded.
7. If nothing was still found then the cell is ‘pre-sporulation/stage I’.
8. The FM464 frame average was similarly segmented but the mask multiplied by the forespore mask to give the FM spore feature.
9. Both FM and YPet spore feature Major/Minor Axis, Area and Orientation were calculated by fitting the shape to an ellipse function.
10. Both were then assigned into 2 shape categories based on the aspect ratio, > 1.2 – ‘septa’, otherwise ‘filled’ structure. These correspond to fluorescence only at the linearly extended septa or distributed about the forespore in a rounder shape.
11. If the FM segmentation was ‘septa’, the segmentation was morphologically ‘thinned’ and its linear curvature calculated.
12. Stages were then assigned as follows:

Stage I/pre-sporulation: no YPet spore feature detected.
Stage II_i_: ‘septa’ FM and YPet spore features with curvature <1.
Stage II_ii_: ‘septa’ FM and ‘filled’ YPet spore features.
Stage II_iii_: ‘septa’ FM and YPet spore features with curvature >1.
Stage III: ‘filled’ FM and YPet spore features.

To confirm the spore categorization algorithm we tested it on a series of simulated images (Fig S2A). These were generated by integrating a model point spread function (PSF) over a 3D model for the cell and forespore shape and subsequently noising the image with Poisson noise based on real noise characteristics of our microscope (27). The cell membrane was modelled as a hollow cylinder, capped with hemisphere shells at either end with 1 pixel thick walls. Stage II_i_ septa were modelled as cell width disks while stage II_iii_ septa were modelled as hemispherical shells. Released SpoIIE in stage II_ii_ was modelled as a hemispherical shell capped by a disk while in stage III, it was modelled as a spherical shell. The relevant features for 100 cells in each stage were simulated in the ‘YPet’ and ‘FM4-64’ channels and run through the categorization algorithm as if they were real data with no noise, average noise and the most extreme noise observed in the data. Without noise. 100% of cells were correctly identified, dropping to at worst in stage IIii 79% with average noise and in the extreme case, as low as 42%.

We attempted further confirmation using Principal Component Analysis (PCA), an approach typically used to identify specific conformations or orientations in cryo-electron microscopy data. Data, images in this case, can be broken down into a basis set of eigenvectors or eigenimages which when summed in proportion to their eigenvalues, recreate the original dataset. Its use in live cell fluorescence data is challenging due to the high heterogeneity in size, shape and intensity of the images. Thus spore images were all cropped to 16×16 pixels, rotated and aligned and their intensity normalised (Fig. S2C) before a basis set of eigenvectors were calculated by Hotelling’s deflation (60). The distribution of eigenvalues was strongly biased towards the 1^st^ eigenvector (Fig. S2D) however 3D scatter plots of the first 3 eigenvalues did show separation of the data, further confirming our categorization algorithm but not allowing us to categorise spores based on PCA alone.

#### Determining the contribution from out-of-focus SpoIIE-YPet foci

To quantify the contribution from out-of-focus SpoIE-YPet foci (i.e. those not detected during tracking) into the membrane ‘pool’ (i.e. spatially extended membranous regions of fluorescence intensity not detected as distinct foci), we assumed that the number and stoichiometry of detected foci from within the depth of field extended were the same as those without and a uniform distribution. Assuming a depth of field of ~350nm, on the basis of expectations from the numerical aperture of the objective lens and peak emission wavelength, a mean cell width of ~0.9 μm (61) and that the focal plane is exactly on the cell midplane we estimate ~1/4 of the cell membrane lies in the depth of field of the microscope. Thus, to generate indicative estimate for copy number values per cell we extrapolated the total number of summed SpoIIE-YPet in foci by a factor of 4x. For the stage II mother cell (Table S2), the mean total number of molecules in foci per cell is ~32 (Mean foci stoichiometry multiplied by mean number of foci per cell) which multiplied by 4 agrees with the mean copy number of 82±42 to within experimental error. Using the same method on other stages either agrees or over or under estimates implying that there is no measurable diffusive ‘pool’ of SpoIIE i.e. all of the SpoIIE-YPet fluorescence can be accounted for by foci.

#### Simulating the effects of different oligomeric states for SpoIIE on the predicted stoichiometry distribution from Slimfield analysis

To simulate the effects of different oligomeric states of SpoIIE-YPet on the observed stoichiometry distribution from Slimfield image data we calculated the probably of foci overlap (56) in each individual cell using the number of detected spots and the area of the spore feature in that particular cell. This probability was used to generate the distribution of overlaps using a Poisson distribution. The predicted apparent stoichiometry distribution was then generated by convolving the overlap distribution with the intensity distribution of model stoichiometry, *S* (i.e. *S*=2, dimers, *S*=4, tetramers etc.). This intensity distribution was generated from the YPet characteristic intensity distribution (Fig. S3C), re-centred on 2*S*, width scaled to *S*^1/2^*σ, where σ =0.675, the sigma width of Fig. S3C. Finally, each of these modelled cell stoichiometry distributions was averaged over the sporulation stage population to generate the model distribution. The goodness-of-fit *R*^*2*^ values for the tetramer model were found to be the highest. (Fig. 3 and S5)

#### Modeling the frictional drag on SpoIIE foci

We modeled the frictional drag coefficient in the cell membrane of SpoIIE foci as that due to a cylinder whose height *h* matches the width of the phospholipid bilayer (~3nm) with a radius given by parameter *a*, using a generalized method established previously to characterize the lateral diffusion of transmembrane proteins (35, 36). In brief, the diffusion coefficient *D* is estimated from the Stokes-Einstein relation of *D*=*k*_*B*_*T*/γ, where *k*_*B*_ is the Boltzmann constant and *T* the absolute temperature, and the lateral viscous drag γ is given by:
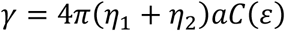
where *η*_1_ and *η*_2_ are the dynamic viscosity values either side of the membrane, which we assume here are approximately the same at *η*_c_ the cytoplasmic viscosity. *C* is a function of Є=2*a*η_c_/*h*η_m_ where *η*_m_ is the dynamic viscosity in the membrane itself. Since *η*_m_ is typically 2-3 orders of magnitude larger than *η*_c_ (34) then *ε* is sufficiently small to use an approximation for *C* of:
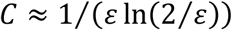

We used these formulations to generate a look-up table between *D* and *a* for the vegetative cell membrane in the mother cell, assuming *η*_m_≈600 cP, and the emerging forespore cell membrane, assuming *η*_m_≈1,000 cP, assuming *η*_c_≈1 cP throughout (Fig. 5C) (37). We estimated a consensus value for *D* in the mother cell from the population of unweighted mean *D* values determined from all cell stages I-III (Table S2) of 1.05 ± 0.06 μ^2^m/s (±SEM, number of stages n=5), and similarly estimated a consensus *D* value for the low mobility sporulation stages II_i_ and II_iii_ of 0.47 ± 0.04 μ^2^m/s (number of stages n=2) and a consensus *D* value for the high mobility sporulation stages II_ii_ and III of 0.76 ± 0.05 μ^2^m/s (number of stages n=2). We then extrapolated these consensus values and SEM error estimates using the vegetative and forespore cell membrane look-up tables to determine corresponding mean values and ±SEM ranges for *a*.

#### Stoichiometry *vs*. localisation

To compare foci stoichiometry as a function of location in the forespore a simplified, normalised 1D coordinate was used. This was based on the generous forespore segmentation which extends from the mother cell side of the septa through to the outer edge of the cell pole. There was also significant variation in the size of this segmentation between cells. Thus a normalised coordinate was used, 0-1 from the two most extreme points of the forespore. This implied that on average both the septa and cell poles lie within the most extreme points of the predicted cell outline segmentation.

## Supplementary Movie Legends

Supplementary Movie 1: Single-molecule movie of SpoIIE-YPet cells in pre-septation. YPet florescence in green, intensity displayed between 345 and 5000 counts. Cell and forespore segmentation indicated as white dots. Tracked foci trajectories indicated as white lines. Time in milliseconds is indicated. Scale bar 1μm.

Supplementary Movie 2: Single-molecule movie of SpoIIE-YPet cells in septation (Stage II_i_ of sporulation). YPet florescence in green, intensity displayed between 345 and 5000 counts. Cell and forespore segmentation indicated as white dots. Tracked foci trajectories indicated as white lines. Time in milliseconds indicated. Scale bar 1μm.

Supplementary Movie 3: Single-molecule movie of SpoIIE-YPet cells in post engulfment (Stage III of sporulation). YPet florescence in green, intensity displayed between 345 and 5000 counts. Cell and forespore segmentation indicated as white dots. Tracked foci trajectories indicated as white lines. Time in milliseconds indicated. Scale bar 1μm.

## Supplementary Figures

**Fig. S1.**
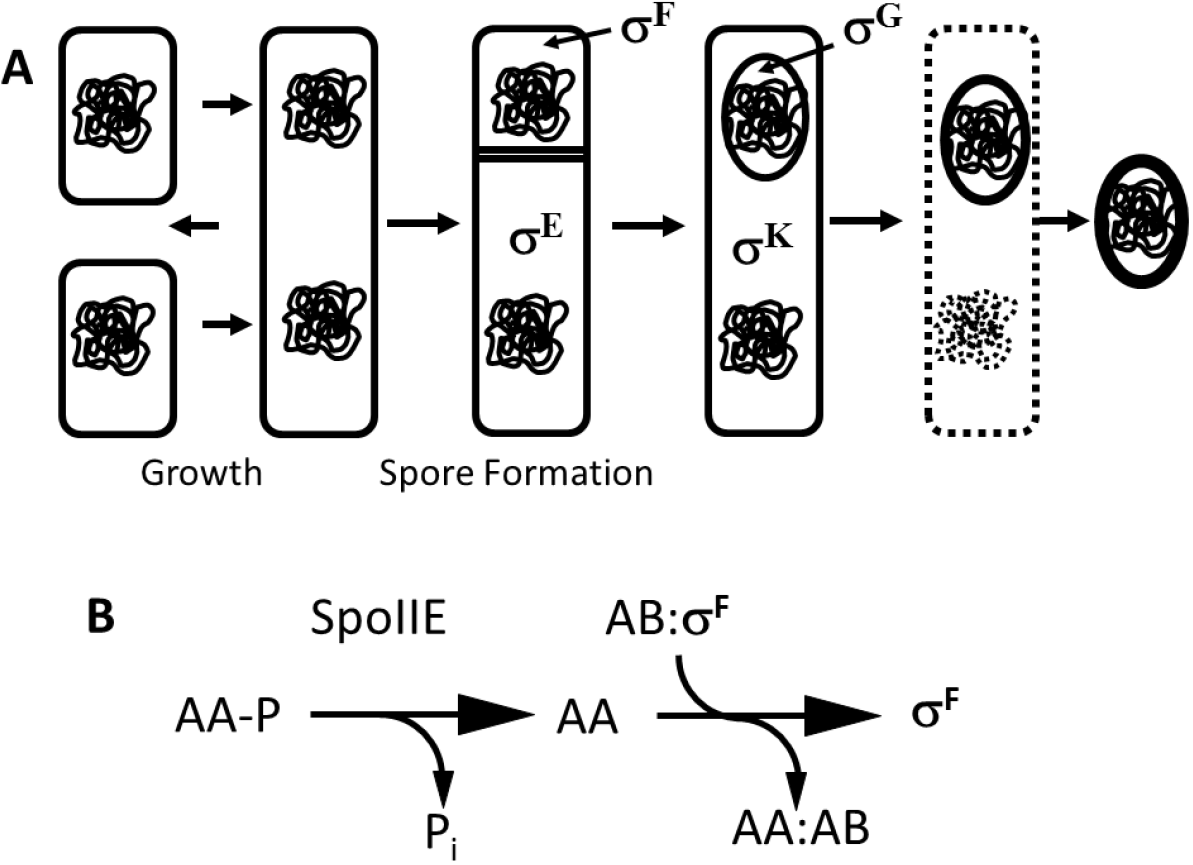
A. Growth and sporulation of *B. subitlis*. During spore formation, the cell divides asymmetrically producing a smaller forespore and a larger mother cell. Compartment and stage specific sigma factors are activated sequentially. The forespore is engulfed by the mother cell before maturing into a resistant spore which is released when the mother cell lyses. B. The SpoIIE phosphatase is the most upstream-acting of three proteins regulating the activity of the first compartment-specific sigma factor, σ^F^. Dephosphorylation allows SpoIIAA (AA) to displace σ^F^ from its complex with an anti-sigma factor (AB) enabling forespore-specific gene expression to be established.

**Fig. S2.**
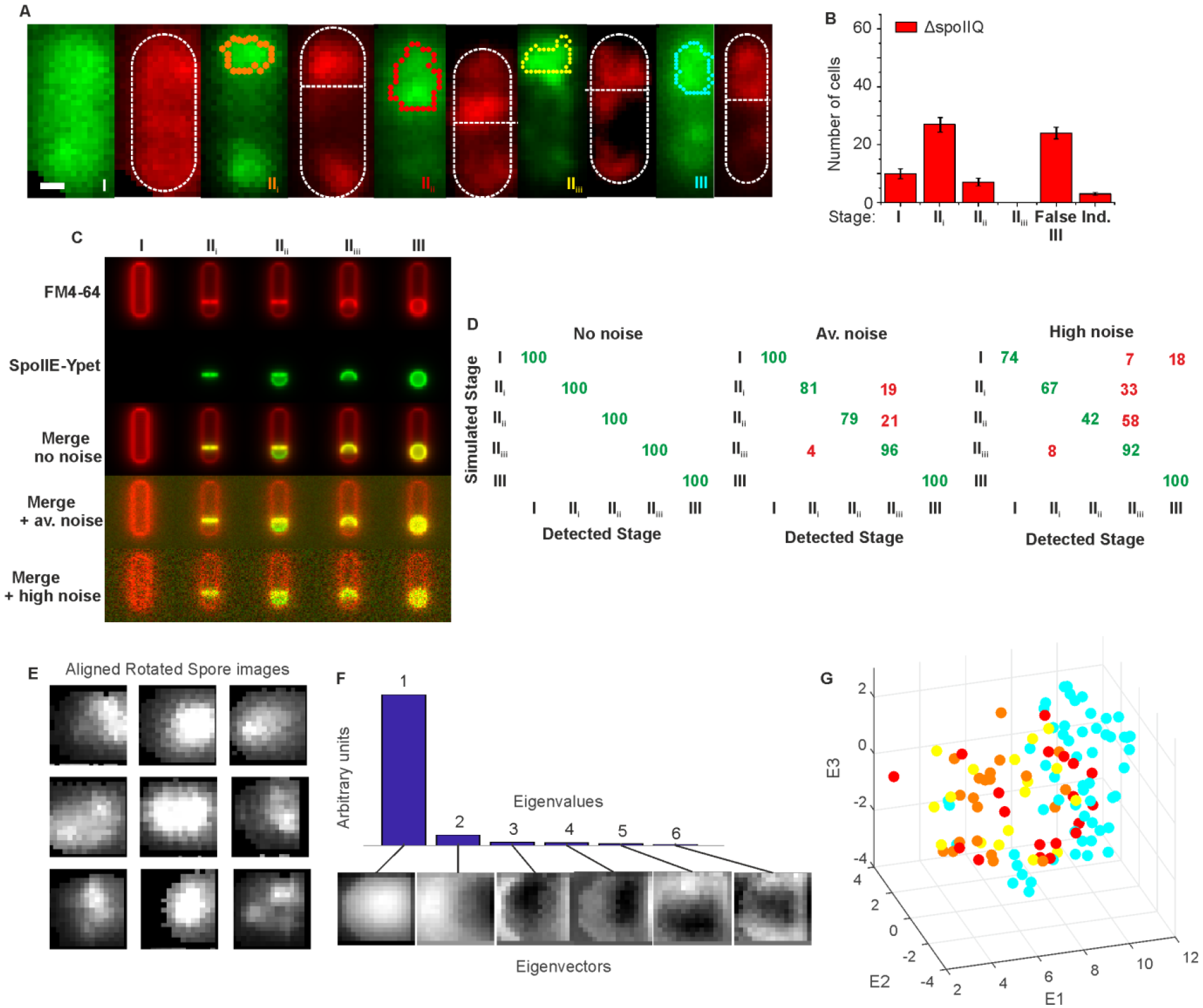
Characterising sporulation stage categorization algorithm. (A) Slimfield 5 frame average images of automatically detected single cells and forespores, categorised by sporulation stages, with detected forespore/septa feature indicated on the SpoIIE image and the cell and forespore boundary indicated in white on the FM4-64 image, scale bar 0.5μm.(B) Frequency of detected cells in each sporulation stage or indeterminate stage (Ind.) for SpoIIE, *ΔspoIIQ* (C) Simulated images of FM4-64 and SpoIIE-YPet at stages I-III without and with representative or high levels of noise. (D) Summary of categorization of 100 simulated images at each stage with the different noise conditions. (E) Aligned, rotated and cropped images of forespores for PCA analysis. (F) Eigenvalues and eigenvectors from PCA analysis. (G) Scatter plot of the relative weights of the first 3 eigenvectors for ~100 cells. The colors represent different stages of cell development as specified in the main text figures.

**Fig. S3.**
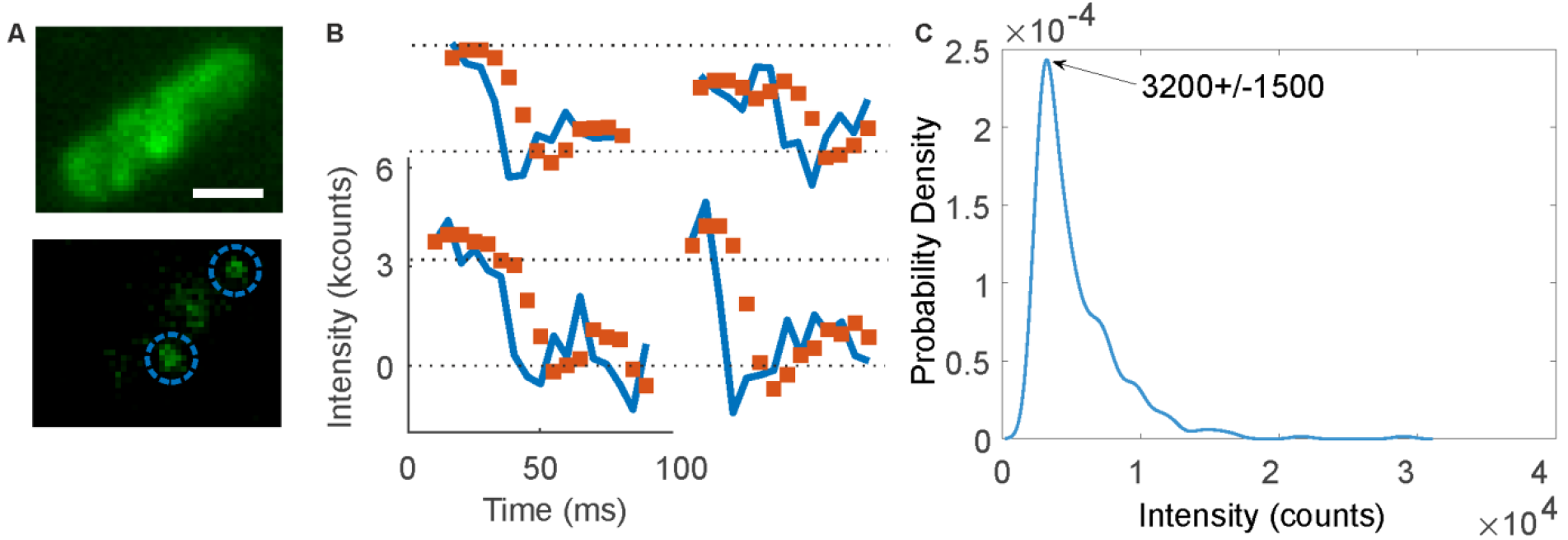
Characteristic *in vivo* intensity of SpoIIE-YPet for stoichiometry determination. (A) Representative SpoIIE slimfield image before photobleaching (above) and towards the end of the photobleach (below) when single SpoIIE-Ypet foci become visible. (B) Overtracked foci intensity as a function of time. Raw data in blue and edge-preserving Chung-Kennedy filtered in red. Steps indicate individual SpoIIE-YPet molecules. (C) Distribution of foci intensity towards the end of the photobleach. Peak value (with +/-HWHM indicated) yields the characteristic SpoIIE-YPet intensity.

**Fig. S4.**
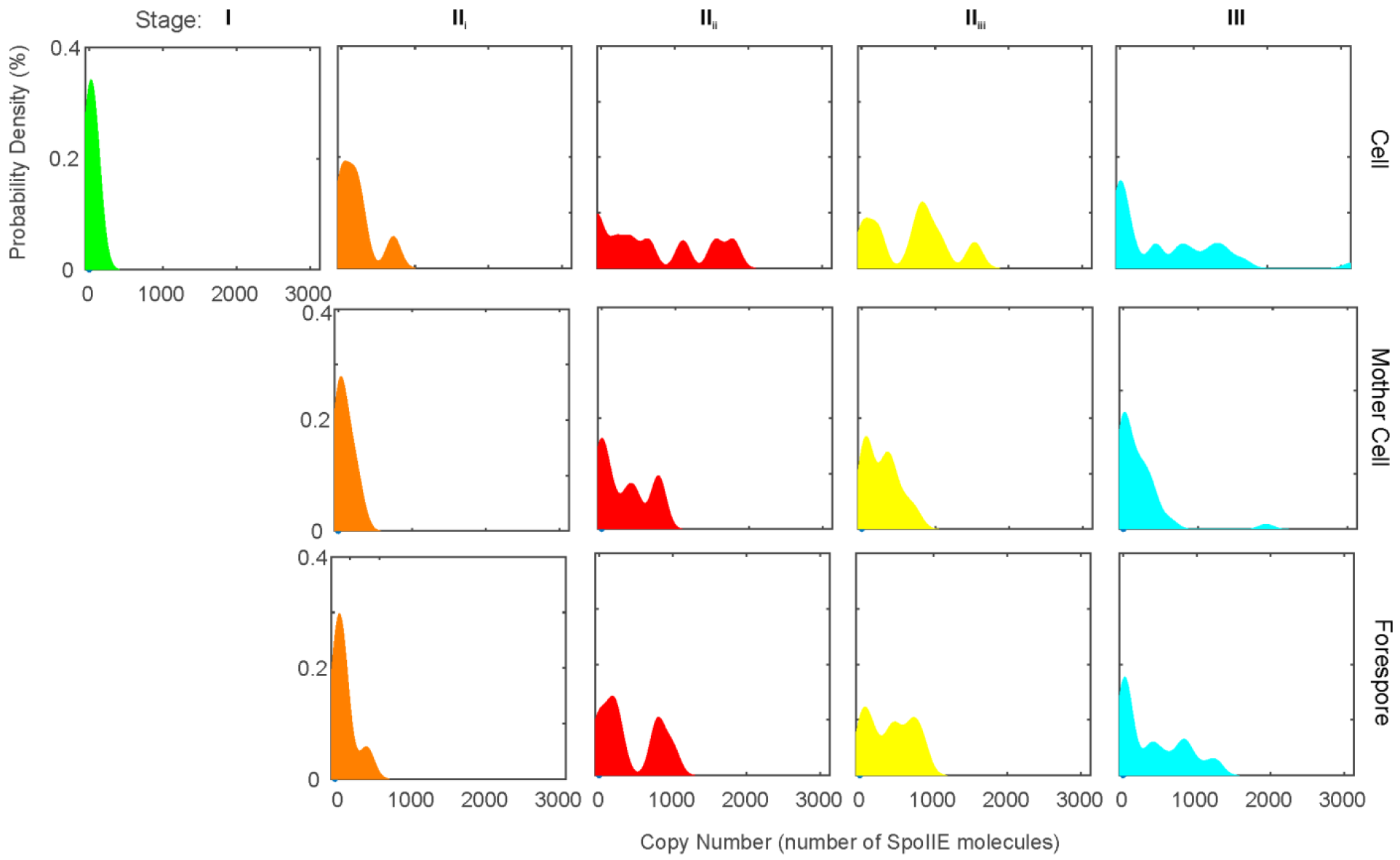
KDE of copy numbers in the whole cell, mother cell and forespore across stages I-III.

**Fig. S5.**
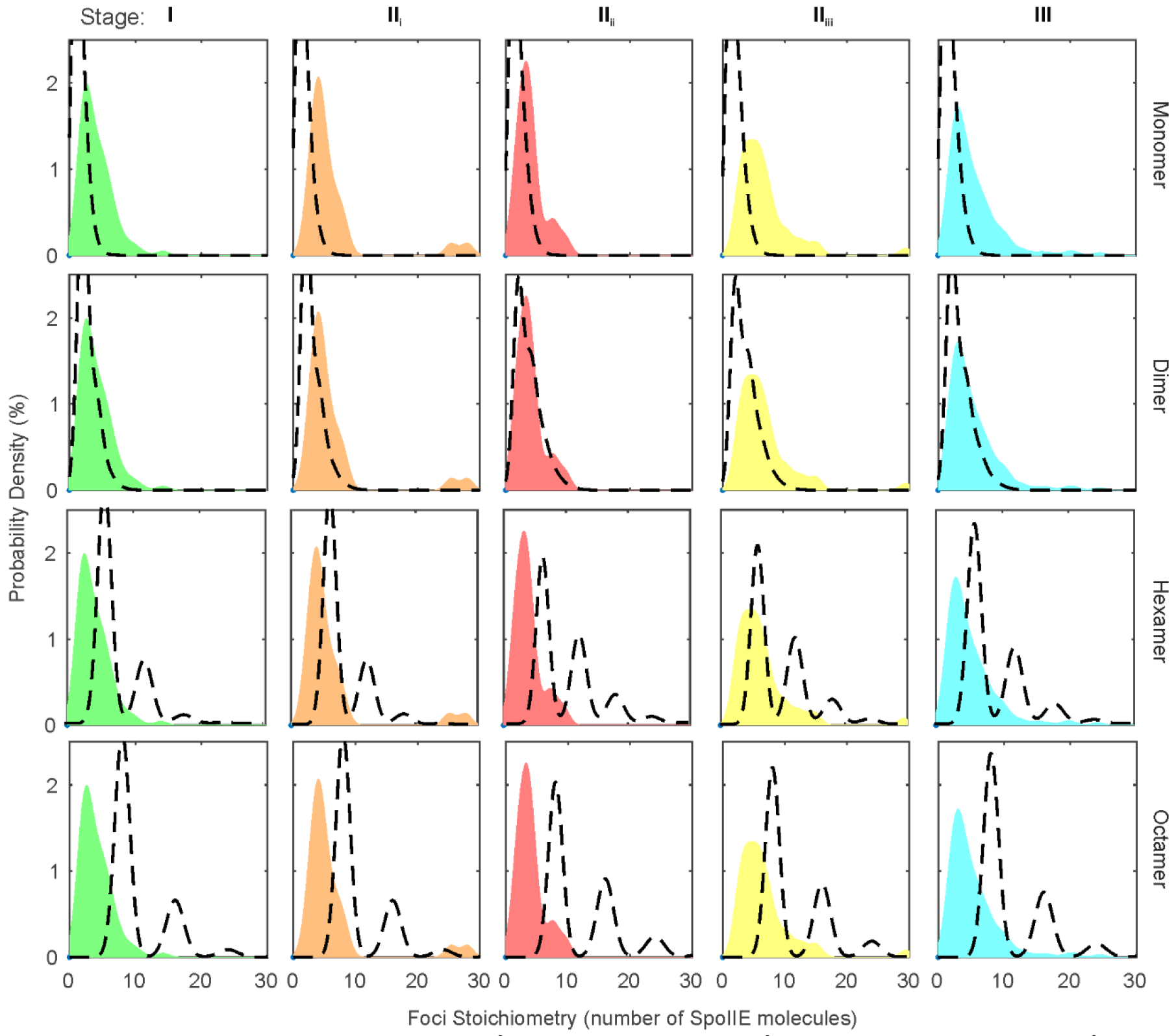
Monomer (goodness-of-fit *R*^*2*^=-5 to -1), dimer (*R*^*2*^=-1.5 to -1), hexamer (*R*^*2*^=-0.5 to 0.7) and octamer (*R*^*2*^ =-1 to -0.5), stoichiometry models compared to mother cell stoichiometry KDE.

**Fig. S6.**
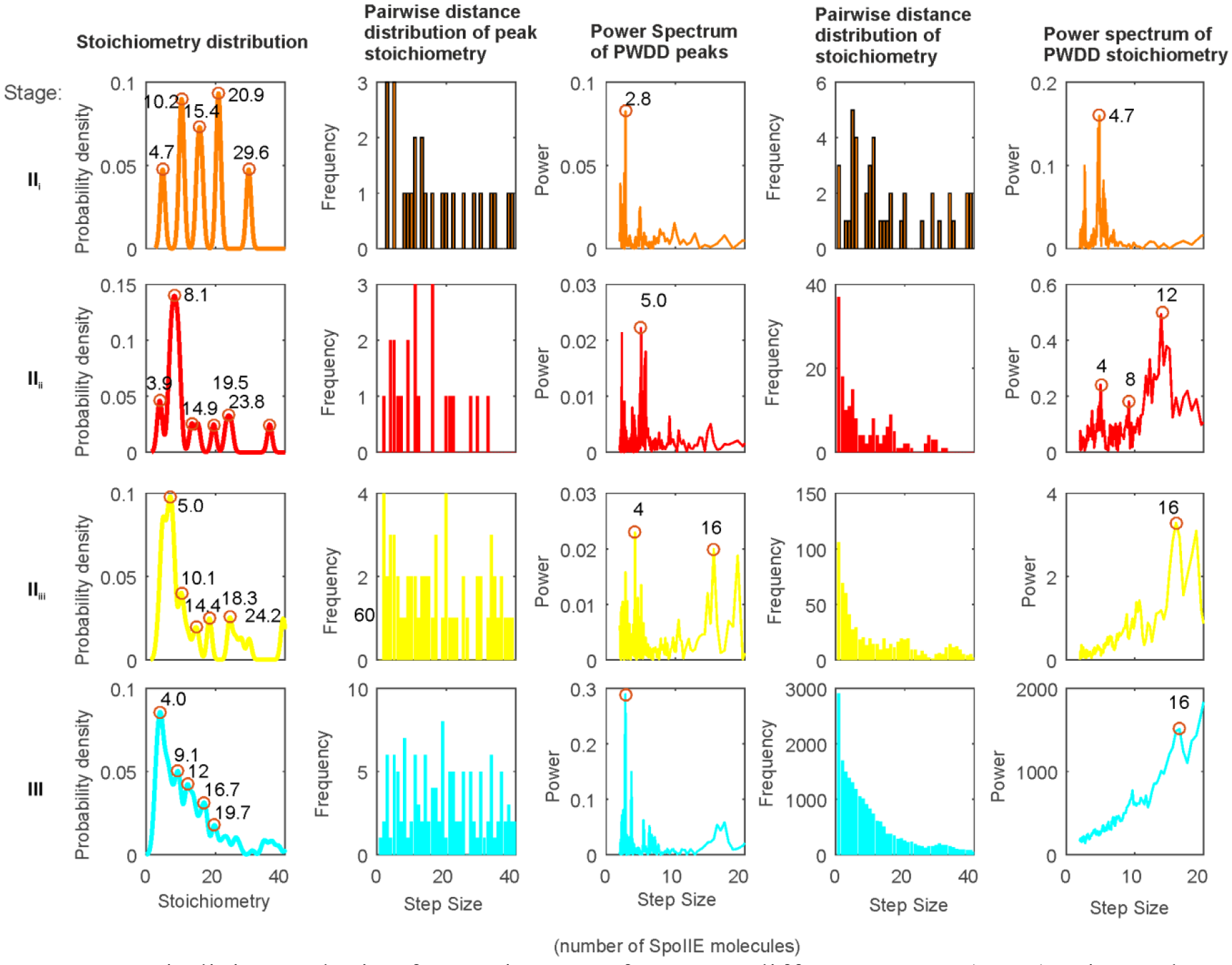
Periodicity analysis of spots in spore feature at different stages (rows). First column contains KDE with kernel width=0.7 molecules. Peak stoichiometries are labelled and are periodic ~4 molecules. Second column shows the pairwise distance distribution (PWDD) of peak values. 4 and multiples occur more frequently. Third column: the power spectrum by Fourier transform of the peak PWDS showing peaks at ~4. Forth Column: PWDS of stoichiometry. Fifth column: Power spectrum of stoichiometry PWDS showing peaks at 4 or harmonics.

**Fig. S7.**
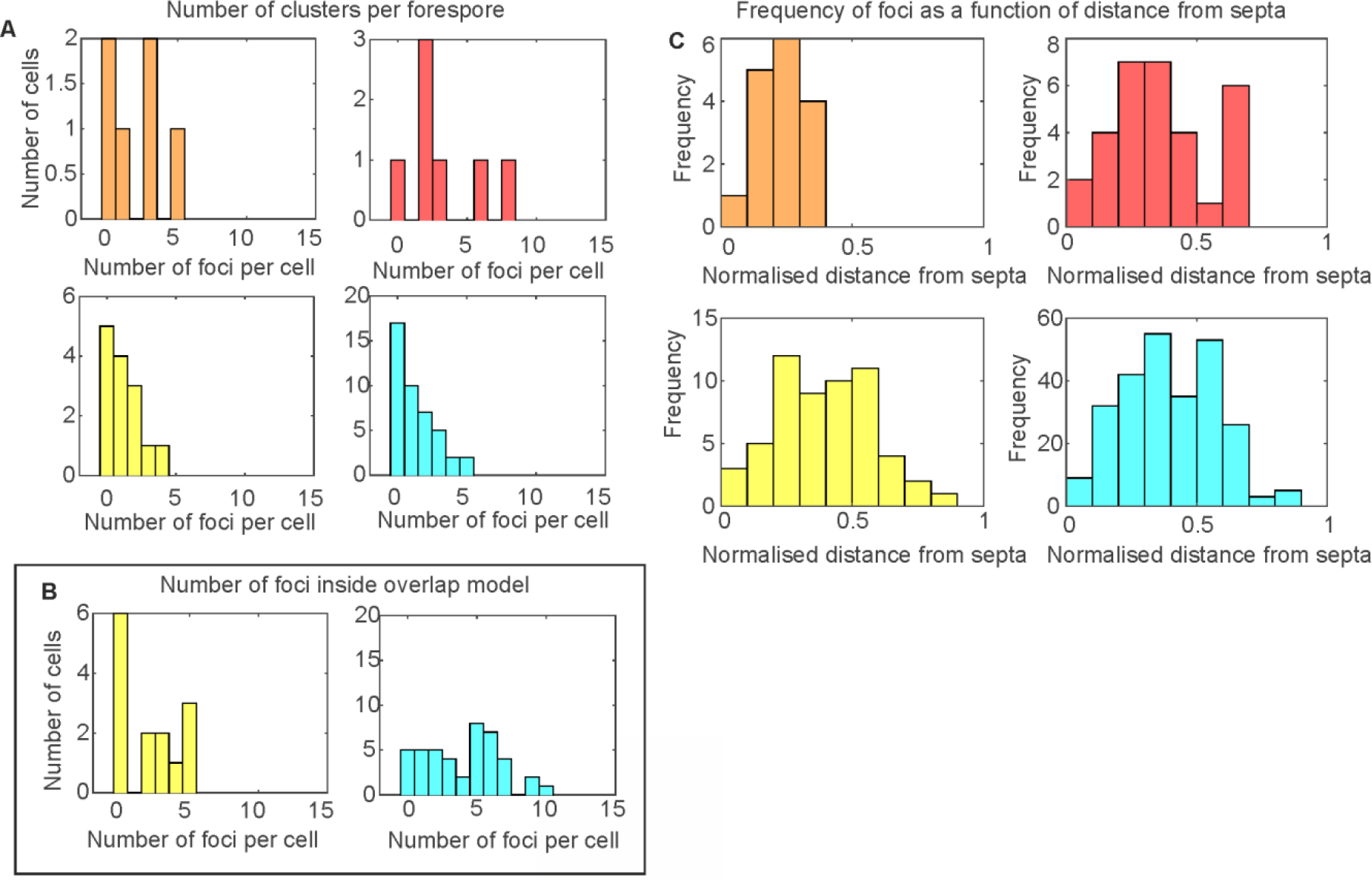
Number of foci found per cell outside (A) and inside (B) the overlapping tetramer model. Foci outside the model and higher stoichiometry SpoIIE clusters. (C) Frequency of foci as a function of normalised distance from septa.

**Fig. S8.**
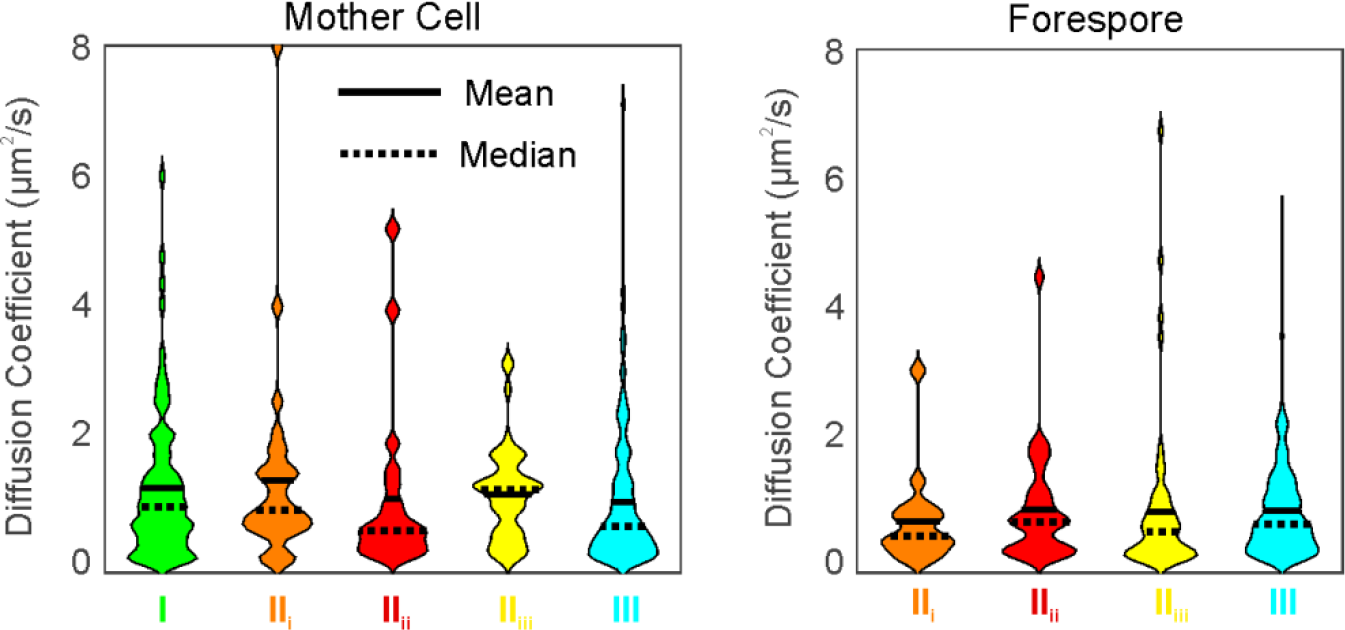
SPoIIE-YPet foci diffusion coefficient distributions rendered as violin plots.

**Fig. S9.**
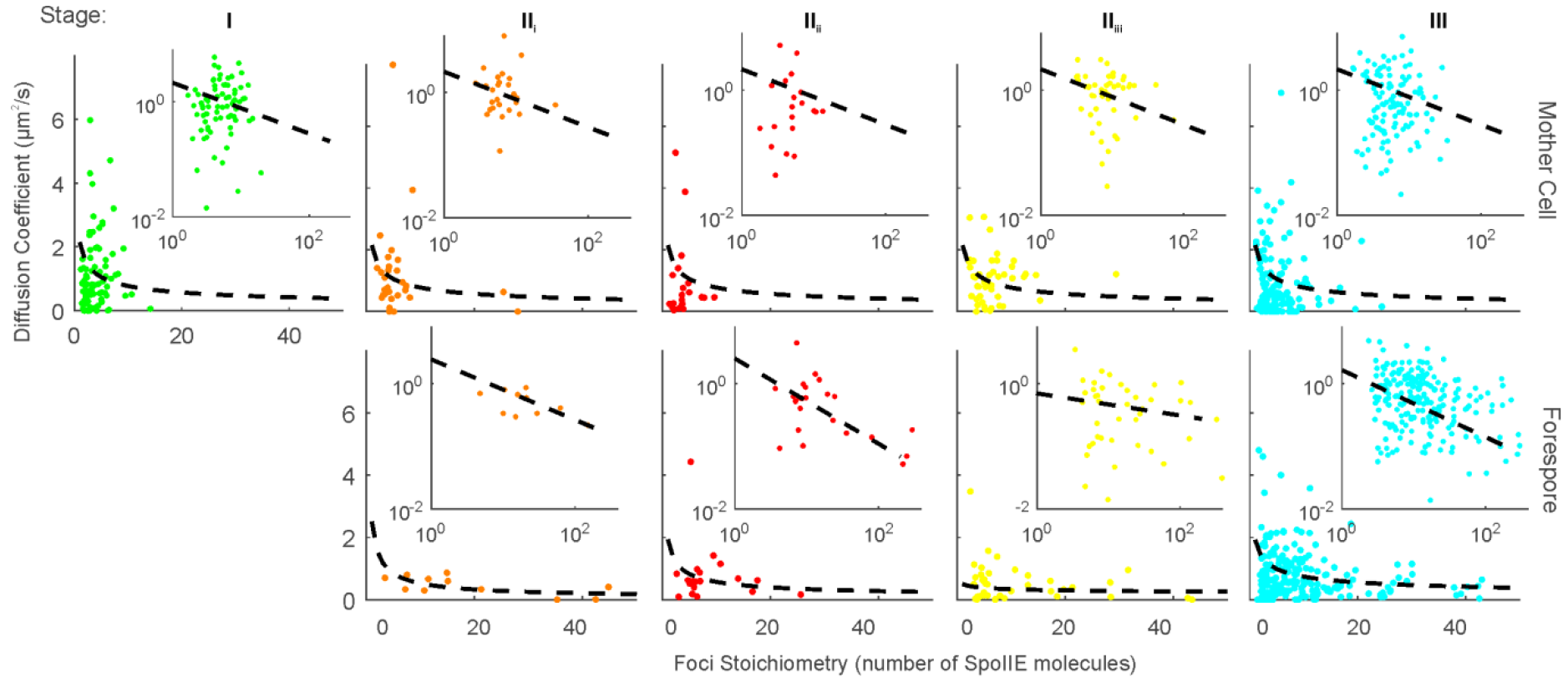
Diffusion coefficient as a function of stoichiometry for tracked SpoIIE-YPet foci for each stage with power law fit overlaid as dotted line. Log-log insert. Each stage fits the same parameters within error.

**Fig. S10.**
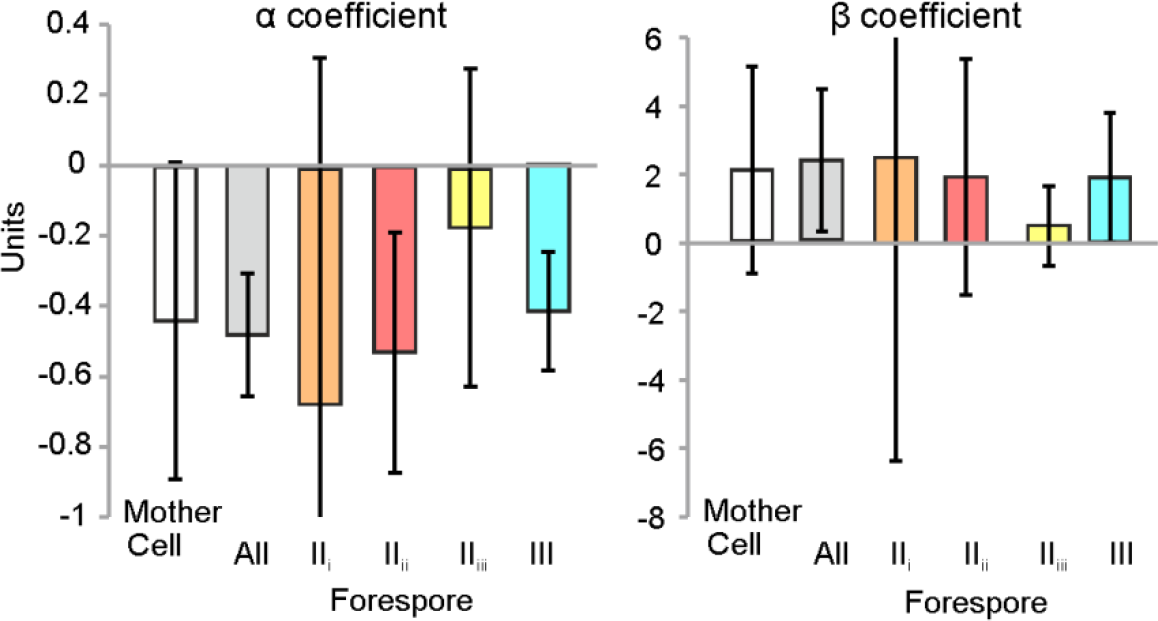
Power law fit coefficients for diffusion coefficient, *D*, as a function of stoichiometry, *S*, *D*=*βS*^*α*^. Each stage fits the same parameters within error.

**Fig. S11.**
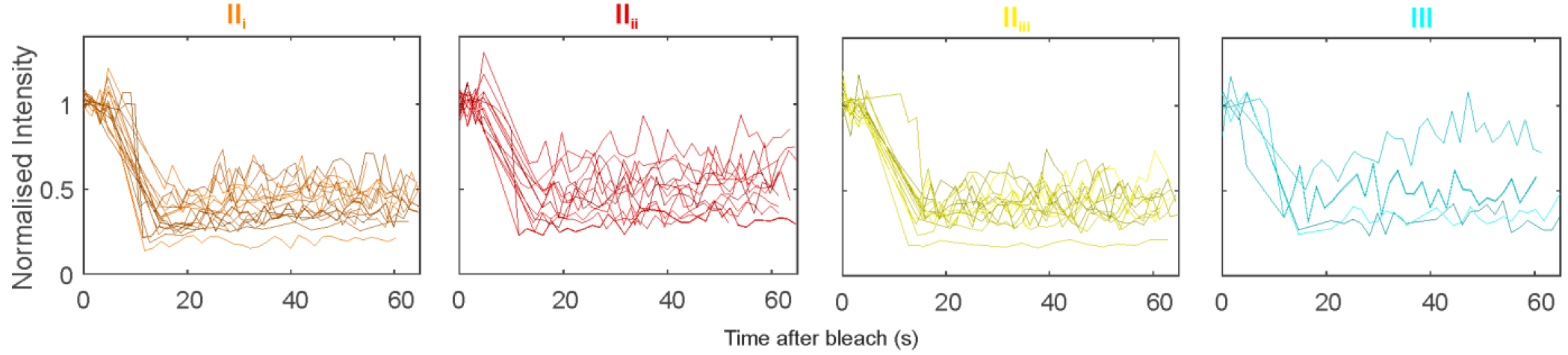
Overlaid plot of all individual cell of normalised fluorescence recovery for each stage.

